# OptoProfilin: A Single Component Biosensor of Applied Cellular Stress

**DOI:** 10.1101/2023.10.04.560945

**Authors:** Noah Mann, Jahiem Hill, Kenneth Wang, Robert M. Hughes

## Abstract

The actin cytoskeleton is a biosensor of cellular stress and a potential prognosticator of human disease. In particular, aberrant cytoskeletal structures such as cofilin-actin rods and stress granules formed in response to energetic and oxidative stress are closely linked to neurodegenerative diseases such as Alzheimer’s, Parkinson’s, and ALS. Whether these cytoskeletal phenomena can be harnessed for the development of biosensors for cytoskeletal dysfunction and, by extension, neurodegenerative disease progression, remains an open question. In this work, we describe the design and development of an optogenetic iteration of profilin, an actin monomer binding protein with critical functions in cytoskeletal dynamics. We demonstrate that this optically activated profilin (‘OptoProfilin’) can act as an optically triggered biosensor of applied cellular stress in select immortalized cell lines. Notably, OptoProfilin is a single component biosensor, likely increasing its utility for experimentalists. While a large body of preexisting work closely links profilin activity with cellular stress and neurodegenerative disease, this, to our knowledge, is the first example of profilin as an optogenetic biosensor of stress-induced changes in the cytoskeleton.

## INTRODUCTION

Profilin is an actin-binding monomer that plays diverse roles in cytoskeletal dynamics, as a supplier of actin to growing filaments (Pring et al., 1992), as a mediator of Arp 2/3 mediated actin filament branching (Mullins et al., 1998), and as part of a ‘pacemaker’ system for the regulation of actin filament growth (Funk et al., 2019). In addition, profilin has recently been implicated in both cofilin-actin rod (Munsie and Truant, 2012; Walter et al., 2021) and stress granule assembly (Figley et al., 2014), strengthening its ties to disease associated cytoskeletal dynamics identified in Alzheimer’s Disease (Borovac et al., 2018; Wurz et al., 2022b) and ALS (Read et al., 2023). These growing ties to neurodegenerative disease indicate that profilin could be incorporated into biosensors strategically designed for early disease detection.

In prior work, we investigated the ability of cofilin and actin mutants, coupled to the blue light responsive *A. thaliana* Cry2/CIB optogenetic system, to act as beacons of cofilin-actin formation in cells exposed to oxidative or energetic stress conditions (Salem et al., 2020). The resulting optogenetic system (‘CofActor’) demonstrated that rapid, stress dependent optogenetic clustering of cofilin and actin can be observed well in advance of endogenous cofilin-actin rod formation. This prompted the question of whether other actin-binding proteins, such as profilin, could also exhibit conditional, light activated responses in cell culture when incorporated into the Cry2 system. To investigate this, an optogenetic fusion protein (Cry2.mCherry.Profilin, **Fig. 1**) was created and investigated for its responses to light activation and cellular stress. Interestingly, when coupled with Cry2-mediated oligomerization, optogenetic profilin displays a condition-dependent light activated response that can be readily observed via fluorescence microscopy. This response is comprised of distinct subcellular localization patterns that differ between stressed and unstressed conditions in cell culture, enabling discrimination of stressed- and unstressed-cell populations. Notably, this Cry2-Profilin conjugate (‘OptoProfilin’) acts as a stand-alone, single component biosensor (e.g., does not require the presence of a Cry2 binding protein such as CIB, distinguishing it from ‘CofActor’), which may increase its general utility in cell-based applications. Opto-biochemical analysis of the observed OptoProfilin response implicates the actin regulatory protein VASP as a critical binding partner of Profilin, and stress-associated changes in VASP localization as the basis for the OptoProfilin stress response. This report demonstrates the application of OptoProfilin in various cell lines and under various stress conditions, characterizing its ability to report on multiple types of stress inputs (energetic, oxidative, osmotic, etc.) and the contribution of the endogenous biochemistries of different immortalized cell lines to OptoProfilin function.

**FIGURE 1.**
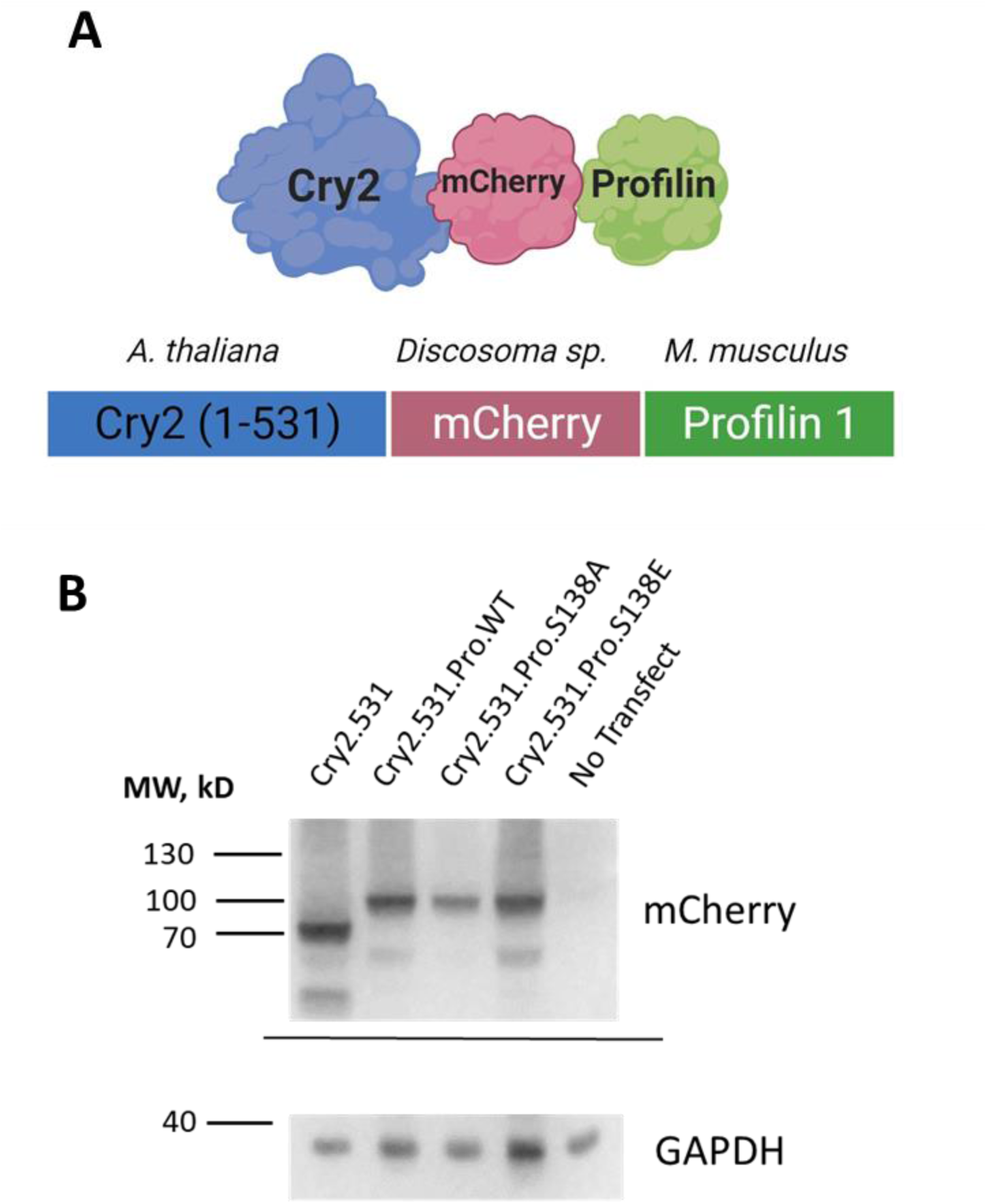
OptoProfilin Construct Design and Expression. A. OptoProfilin is comprised of *A. thaliana* Cryptochrome 2 (aa 1 – 531), *Discosoma sp*. mCherry, and *M. musculus* Profilin 1. *Image created with BioRender.* B. Western blot of lysates from HEK 293T cells transiently transfected with Cry2 (1-531) and OptoProfilin (Cry2.mCh.Profilin) constructs. The molecular weight of the Cry2.mCh.Profilin fusion protein is 103 kD.

## RESULTS AND DISCUSSION

The previously reported CofActor (Cofilin Actin Optically Responsive) system, in which a Cry2.Cofilin S3E fusion and a betaActin.CIB fusion protein form clusters in response to the combination of light and applied energetic or oxidative stress, responds stimuli that also induce native cofilin-actin rods in cells (Salem et al., 2020). We subsequently speculated that other actin binding proteins, such as profilin, might be used in place of cofilin within a similar optogenetic framework. To test this, a Cry2 - Profilin fusion construct (‘OptoProfilin’) was generated (**Fig. 1**) and investigated for its response to either light or a combination of light and energetic stress in HeLa cells. Surprisingly, light activation of the profilin construct in the absence of a CIB binding partner was sufficient to induce a subcellular localization response (**Fig. 2; Supporting Movie 1**), with an apparent targeting of focal adhesions (**Fig. 2A**). To demonstrate that the observed localization pattern was consistent with that of focal adhesions, immunostaining for Paxillin, a focal adhesion localized protein, was performed (**Fig. 2B-C**), demonstrating that optically activated OptoProfilin is co-localized with focal adhesions. We also tested the response of OptoProfilin when exposed to energetic stress (ATP-depletion) conditions. ATP depletion shifted the light-induced localization from focal adhesions to bright, punctate protein clusters (**Fig. 2A**, bottom row; **Supporting Movie 2**). This stress-associated response, while morphologically identical to that observed with CofActor, does not require the presence of an Actin-CIB fusion protein. Thus, OptoProfilin can function as a single component reporter of applied cellular stress.

**FIGURE 2.**
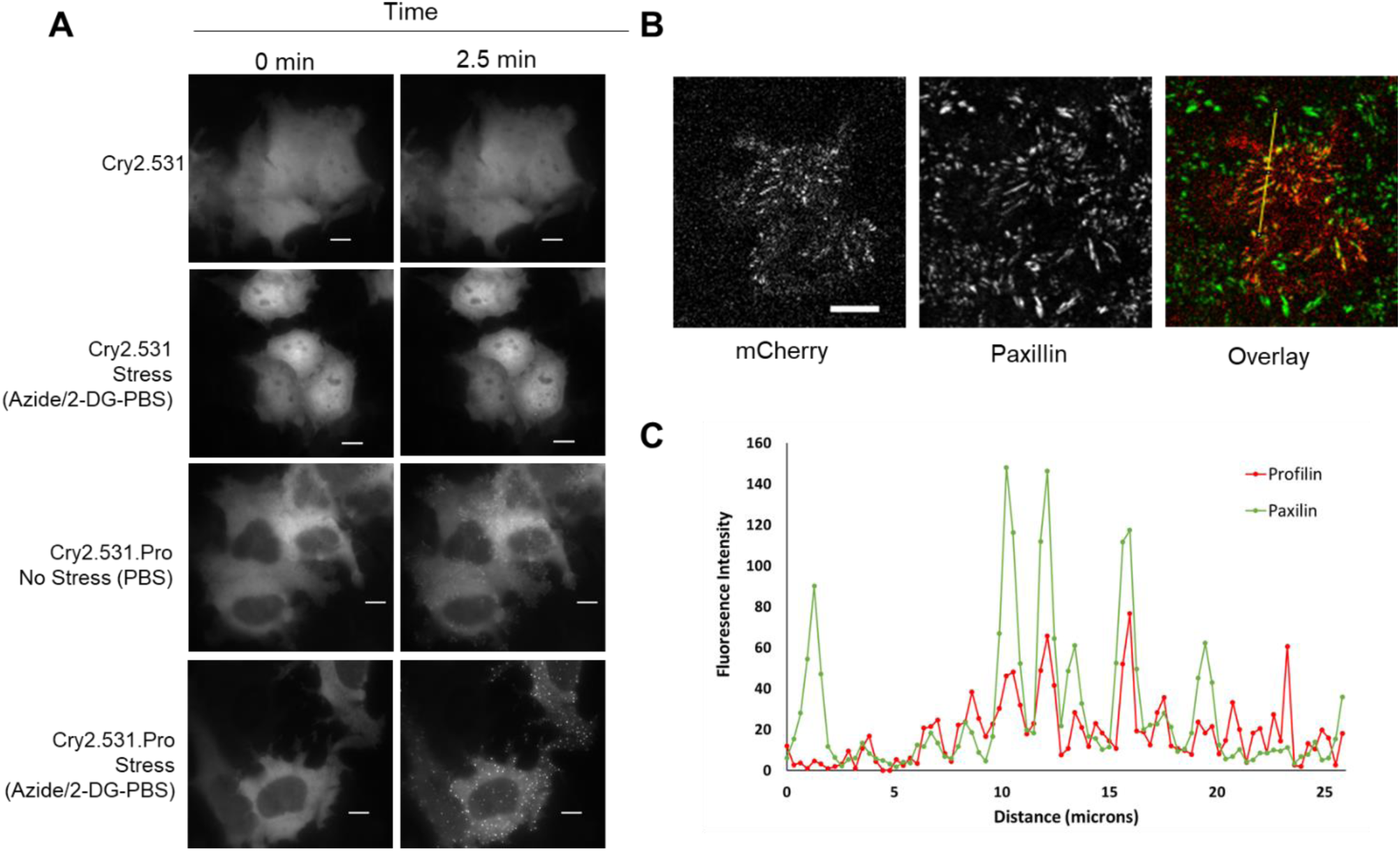
Condition-dependent recruitment of OptoProfilin in HeLa cells. A. Cry2(1-531) exhibits minimal light activated response under buffer-only (top row) or ATP-depletion (second row; 10 mM NaN3/6 mM 2-DG) conditions. Upon light exposure, OptoProfilin recruits to focal adhesions under buffer-only conditions (third row) and clusters under ATP-depletion conditions (bottom row). B. Confirmation of focal adhesion recruitment by immunofluorescence of light-exposed and fixed OptoProfilin-expresssion HeLa cells (mCherry, red; Alexa 488/α-Paxillin, green). Scale bars = 10 microns. C. Overlay of fluorescence intensity of mCherry (OptoProfilin) and Alexa 488 (Paxillin) at region indicated by yellow line in panel B.

Subsequent inquiries were directed towards characterizing the biochemical underpinnings of OptoProfilin’s focal-adhesion targeting properties. Whereas the CofActor system was dependent on a stress-induced cofilin-actin interaction, the selective recruitment of Profilin to focal adhesions, in the absence of the Cry2 binding partner CIB, pointed to an alternate binding mechanism. Previously reported investigations of profilin binding to focal adhesions implicated VASP, an actin-regulatory protein that is typically associated with focal adhesions, as a primary mediator of profilin-focal adhesion interactions (Gau et al., 2019), with profilin residue S138 as the key residue driving VASP profilin-interactions and phosphorylation of profilin S138 as a key inhibitor of VASP-profilin binding. Following literature precedent (Gau et al., 2019), we generated both phospho-mimic (S138E) and constitutively active (S138A) mutations of the OptoProfilin. Consistent with previously reported results, Cry2.Pro.S138E did not exhibit light-activated recruitment to focal adhesions, whereas Cry2.Pro.S138A could be recruited to focal adhesions with light (**Fig. 3**). In addition, immunostaining of OptoProfilin mutant S138A recruited to focal adhesions revealed colocalization with endogenous VASP localized to focal adhesions (**Fig. 3C**). Taken together, these results indicate that VASP binding is the key promoter of OptoProfilin recruitment to focal adhesions in HeLa cells.

**FIGURE 3.**
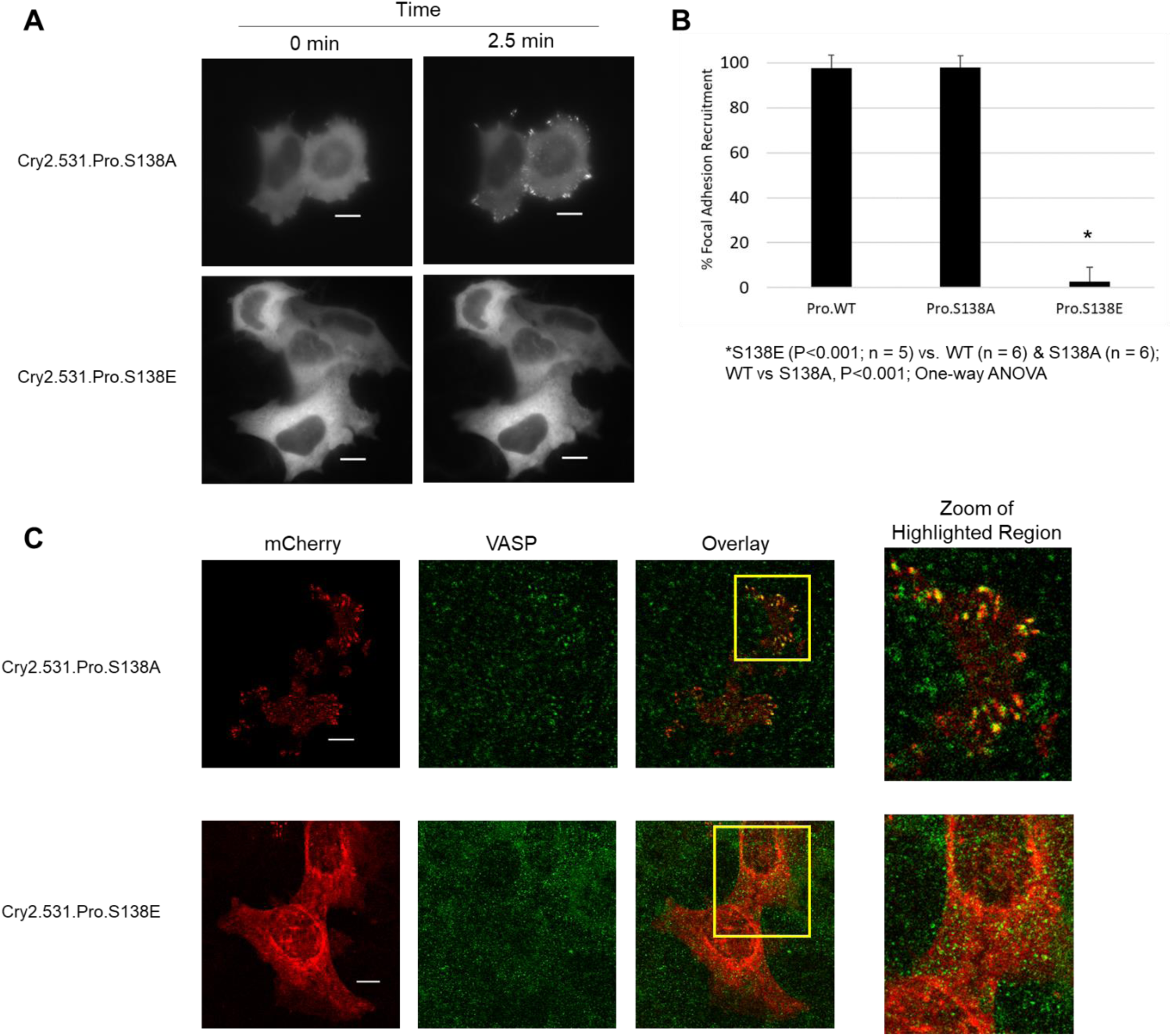
Profilin VASP-binding mutants (S138) display different light-activated recruitment under non-stress conditions. A. OptoProfilin S138A mutant exhibits light-activated recruitment to focal adhesions, but OptoProfilin S138E mutant does not. B. Quantification of percentage of cells exhibiting recruitment of mutants vs. WT OptoProfilin. C. Immunofluorescence of light-activated and fixed OptoProfilin mutants (mCherry, red; Alexa 488/VASP, green) imaged by confocal microscopy. Scale bars = 10 microns.

Next, we investigated the role of Profilin-VASP binding in the ATP depletion-associated optogenetic clustering response. In ATP-depleted HeLa cells, OptoProfilin S138A formed light and stress-induced clusters, while the S138E mutant did not (**Fig. 4 A – B**). In addition, immunostaining of OptoProfilin clusters in ATP-depleted cells demonstrated that the stress-associated clusters also contain endogenous VASP (**Fig. 4 C**). This result indicated that VASP-Profilin interactions are central to both filopodial and cluster recruitment in HeLa cells. To more closely examine the OptoProfilin transition from elongated focal adhesions to bright punctae, we conducted an experiment in which OptoProfilin was activated under non-stress conditions followed by addition of an ATP-depletion inducing stock solution. This experiment revealed that compact VASP-rich clusters emerge directly from elongated focal adhesions (**Fig. 5 A-C; Supporting Movie 3**), generally moving only a short distance from their site of origin. Overall, these experiments demonstrate that the OptoProfilin response is directly correlated with changes in VASP subcellular localization. Nonetheless, the morphology and distribution of stress-associated OptoProfilin clusters closely resembled that of stress granules, which are mRNA-protein condensates that are formed in response to cellular stress and are closely associated with neurodegenerative disease (Protter and Parker, 2016). Furthermore, Profilin1 has been shown to associate with stress granules (Figley et al., 2014). To investigate whether OptoProfilin clusters were similar in composition to stress granules, fixed granules were stained with an RNA-specific stain and immunostained for Ataxin2 (a stress granule core protein) (Li et al., 2013). Staining of these OptoProfilin clusters with an RNA specific dye did not show a strong presence of RNA in the Cry2-Profilin clusters, and Ataxin2 immunostaining was not co-localized with OptoProfilin clusters (**Fig. 6**). This provides evidence that OptoProfilin clusters are compositionally distinct from traditional stress granules.

**FIGURE 4.**
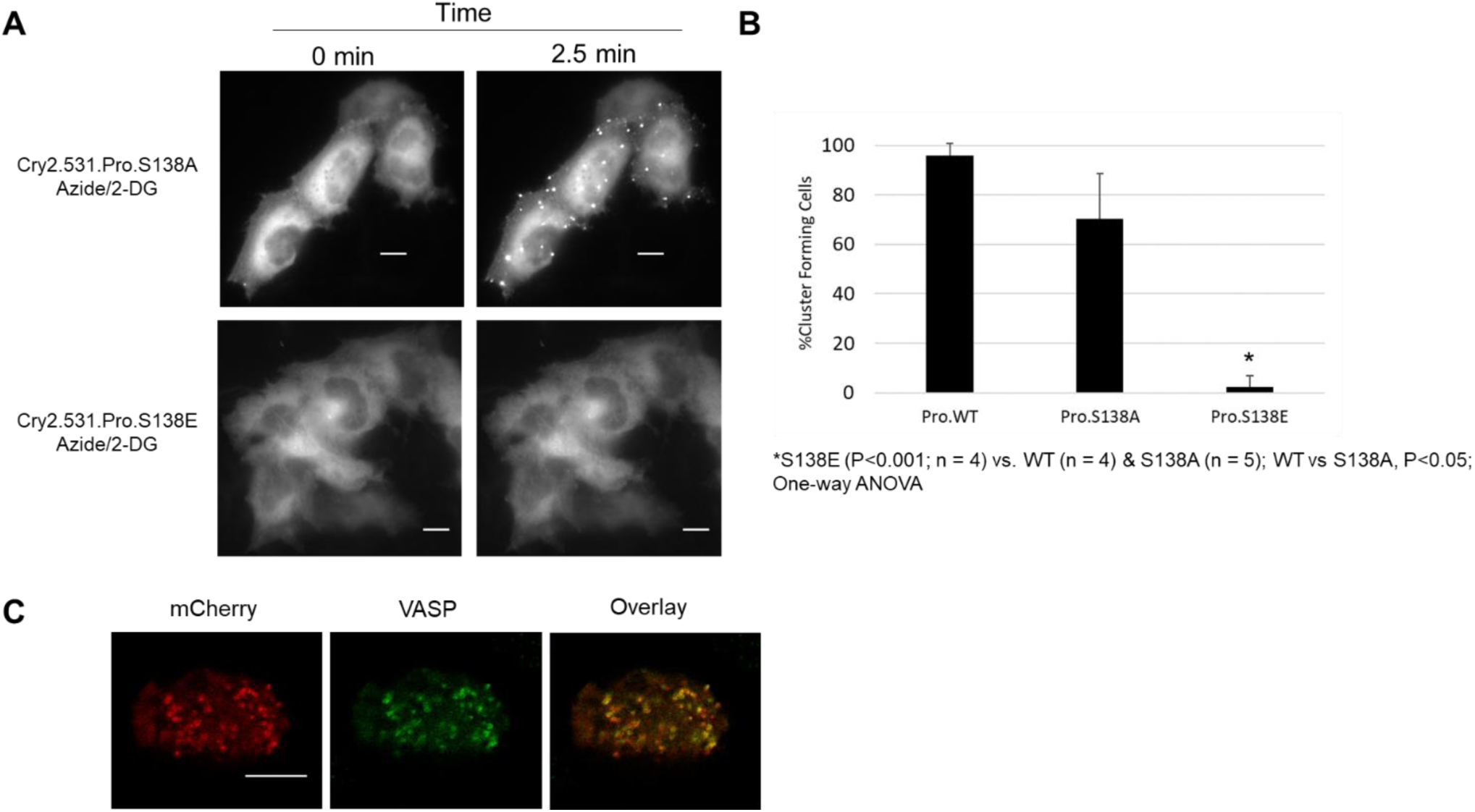
Profilin VASP-binding mutants (S138) display different light-activated recruitment under ATP-depletion conditions. A. OptoProfilin S138A mutant exhibits light-activated recruitment to clusters in ATP-depleted cells, but OptoProfilin S138E mutant does not. B. Quantification of percentage of cells exhibiting cluster recruitment of mutants vs. WT OptoProfilin. C. Immunofluorescence of light-activated and fixed OptoProfilin S138A mutant (mCherry, red; Alexa 488/VASP, green) imaged by confocal microscopy. Scale bars = 10 microns.

**FIGURE 5.**
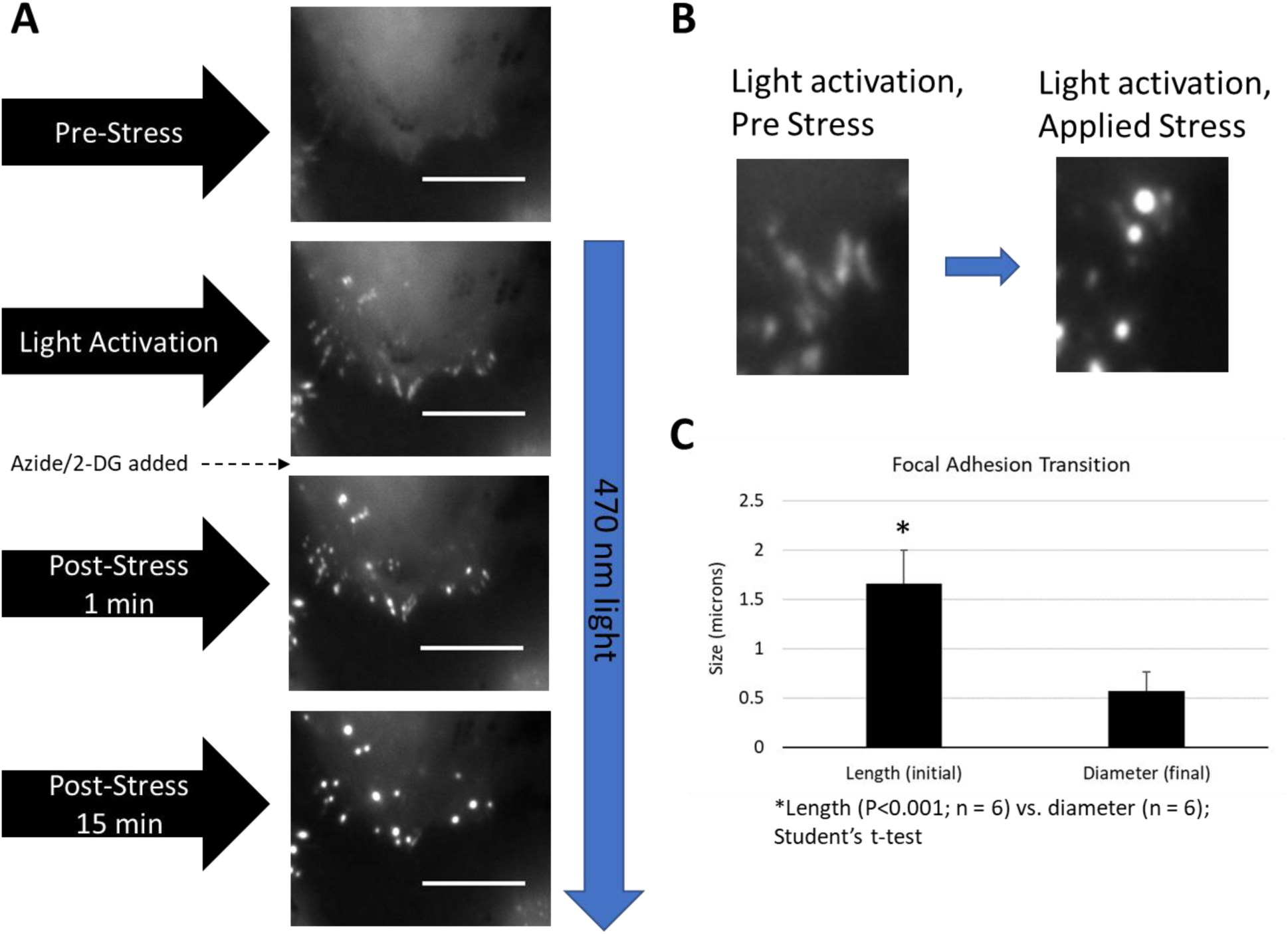
Stress activated, VASP-mediated filopodia to cluster transition. A. OptoProfilin recruitment to elongated filopodia followed by addition of ATP-depletion media (final concentration 10 mM NaN3/6 mM 2-DG) and imaging of subsequent transition to VASP-rich clusters. B. Close up of transition to stress-induced phenotype. C. Quantification of the stress-induced focal adhesion transition observed in panel B. Scale bars = 10 microns.

**FIGURE 6.**
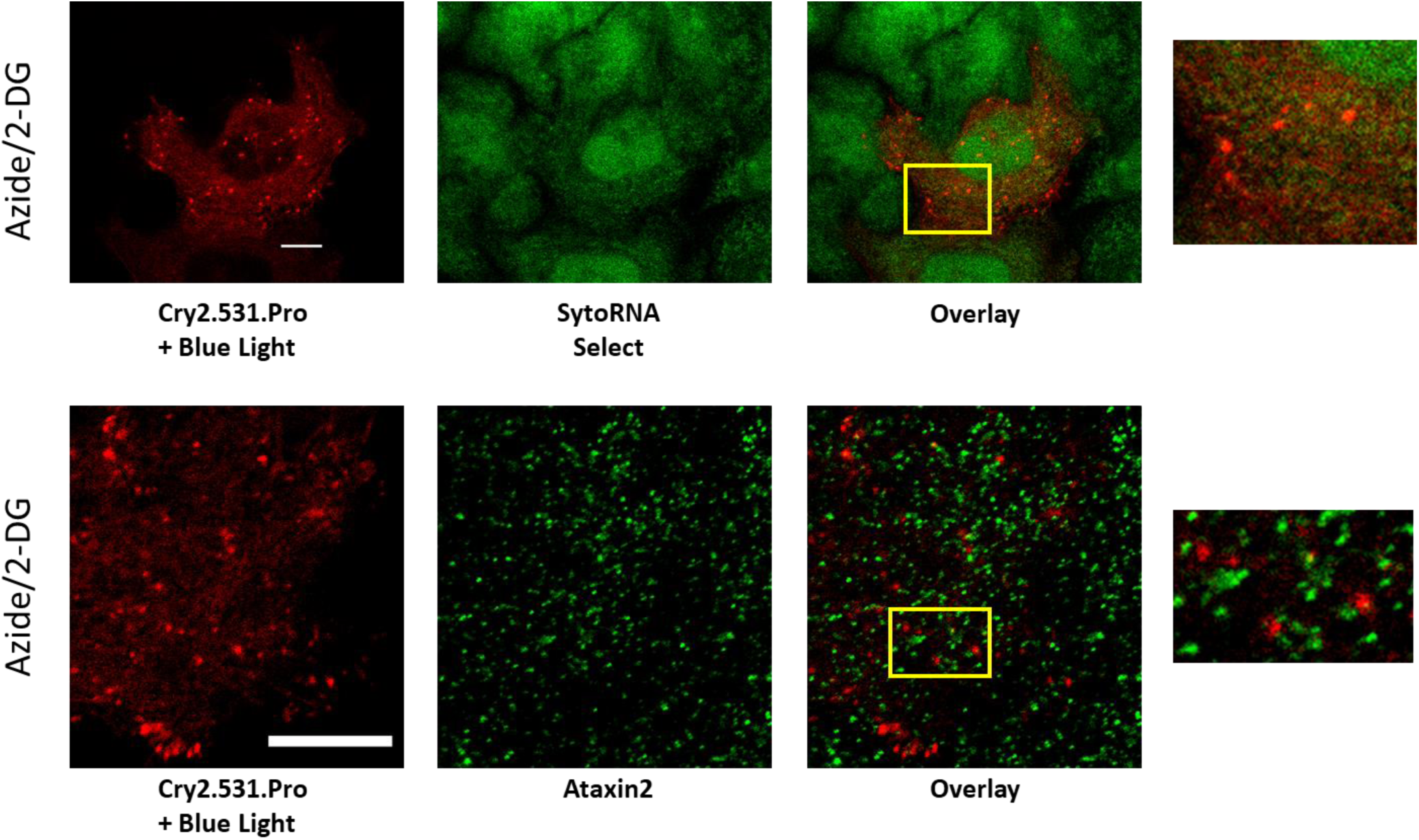
Light and stress activated OptoProfilin cluster contents are not consistent with traditional stress granules. Top: SytoRNA Select stain (green) of light activated ATP-depleted OptoProfilin expressing cells (red). Bottom: Ataxin2 immunofluorescence (green) of light activated ATP-depleted OptoProfilin expressing cells (red). Scale bars = 10 microns.

After investigating OptoProfilin light activated response in ATP-depleted cells, we then turned to investigate other sources of cellular stress. In these experiments, OptoProfilin gave a robust light-activated clustering response to oxidative stress (0.5 mM Sodium Arsenite), osmotic stress (200 mM Sorbitol), and H2O2-induced cellular senescence (200 µM H_2_O_2_ followed by washout and a 72 h post-stress incubation period) (**Fig. 7; Supporting Movie 4**). Heat stress (46 °C, 1h) treatment was the exception, in which OptoProfilin did not form clusters in response to light activation and largely retained the elongated focal adhesion recruitment phenotype. These experiments indicate the presence of a general mechanism enabling the formation of VASP clusters and subsequent OptoProfilin recruitment. Possibilities for this general effect include stressor-induced changes in membrane tension coupled with biochemical changes to VASP and Profilin. While this mechanism is currently under further investigation, common kinase-dependent VASP phosphorylation changes (Ser 157; Ser 239) do not explain the general stress response, as we observed no correlation between trends in VASP phosphorylation at Ser 157 and Ser 239 and various stress treatments (**Fig. 11A-B**). However, it is anticipated that the core components (i.e., VASP) of stress-associated OptoProfilin clusters are the same regardless of stress stimuli. For example, similar to Azide/2-DG-induced clusters, Sodium Arsenite-induced clusters also immunostained for VASP (**Supporting Fig. 1**).

**FIGURE 7.**
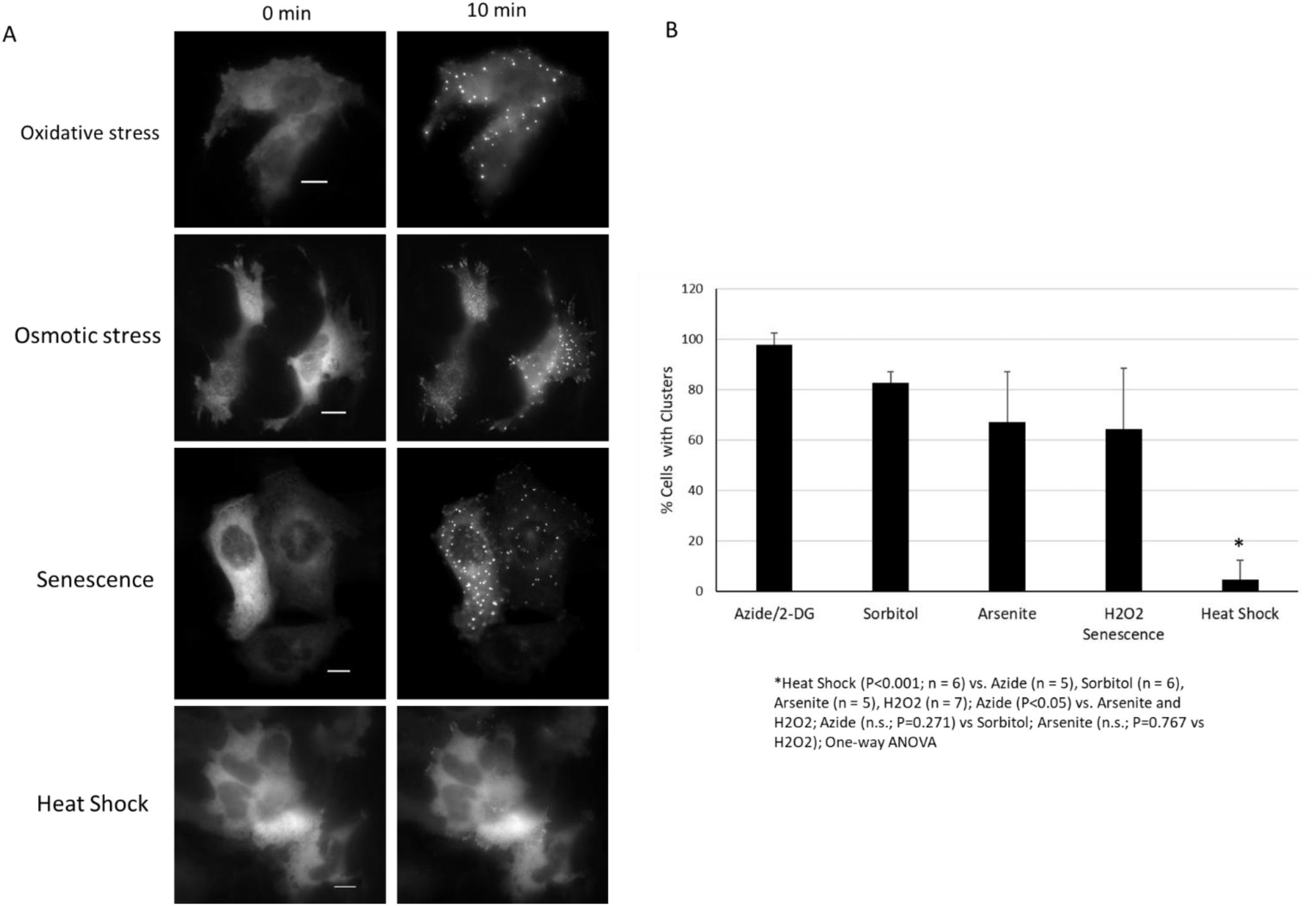
OptoProfilin clustering response to different cellular stress inputs. A. A light-activated clustering response is observed in response to numerous stress inputs, including oxidative stress (0.5 mM Sodium Arsenite, 15 min pre-treatment), Osmotic stress (200 mM Sorbitol, 15 min pre-treatment), Senescence (2 h treatment with 200 µM H_2_O_2_, imaged 72 h post-treatment). Heat shock (46 °C, 1 h) did not induce a clustering response. B. Quantification of experiments described in panel A. Scale bars = 10 microns.

Finally, we investigated the OptoProfilin stress response in cell lines other than HeLa (ex., HEK293T, NIH 3T3, N2a) under ATP depleted conditions. Intriguingly, in cell lines with low endogenous VASP expression (HEK293T and NIH 3T3), the recruitment to elongated focal adhesions was not present in the absence of ATP depletion (**Fig. 8A**, top row; **Fig. 9A**, top row). We attribute this to low VASP expression in these cell lines (**Fig. 8C**), which has also been routinely demonstrated in antibody product literature (Bio-Rad, 2023). However, upon ATP depletion of these cell lines, stress-associated OptoProfilin clusters were readily apparent in both (**Fig. 8 A-B**; **Fig. 9 A-B; Supporting Movie 5**). Generally, these results suggest that application of OptoProfilin as a stress sensor in relatively VASP-deficient cell lines may have lower background signal due to a higher stress response threshold. By contrast, the OptoProfilin response in N2a cells was light-but not stress-gated (**Fig. 10**). Possible origins of the lack of stress-vs. non-stress discrimination in this cell line could be high levels of VASP expression, anomalous VASP localization, or a combination thereof. For example, in the cell lines tested in this study, VASP expression in N2a cells was approximately 1.6-fold higher than that of HeLa cells, and 3-fold higher than that observed in HEK 293 or NIH 3T3 cell lines (**Fig. 11C**).

**FIGURE 8.**
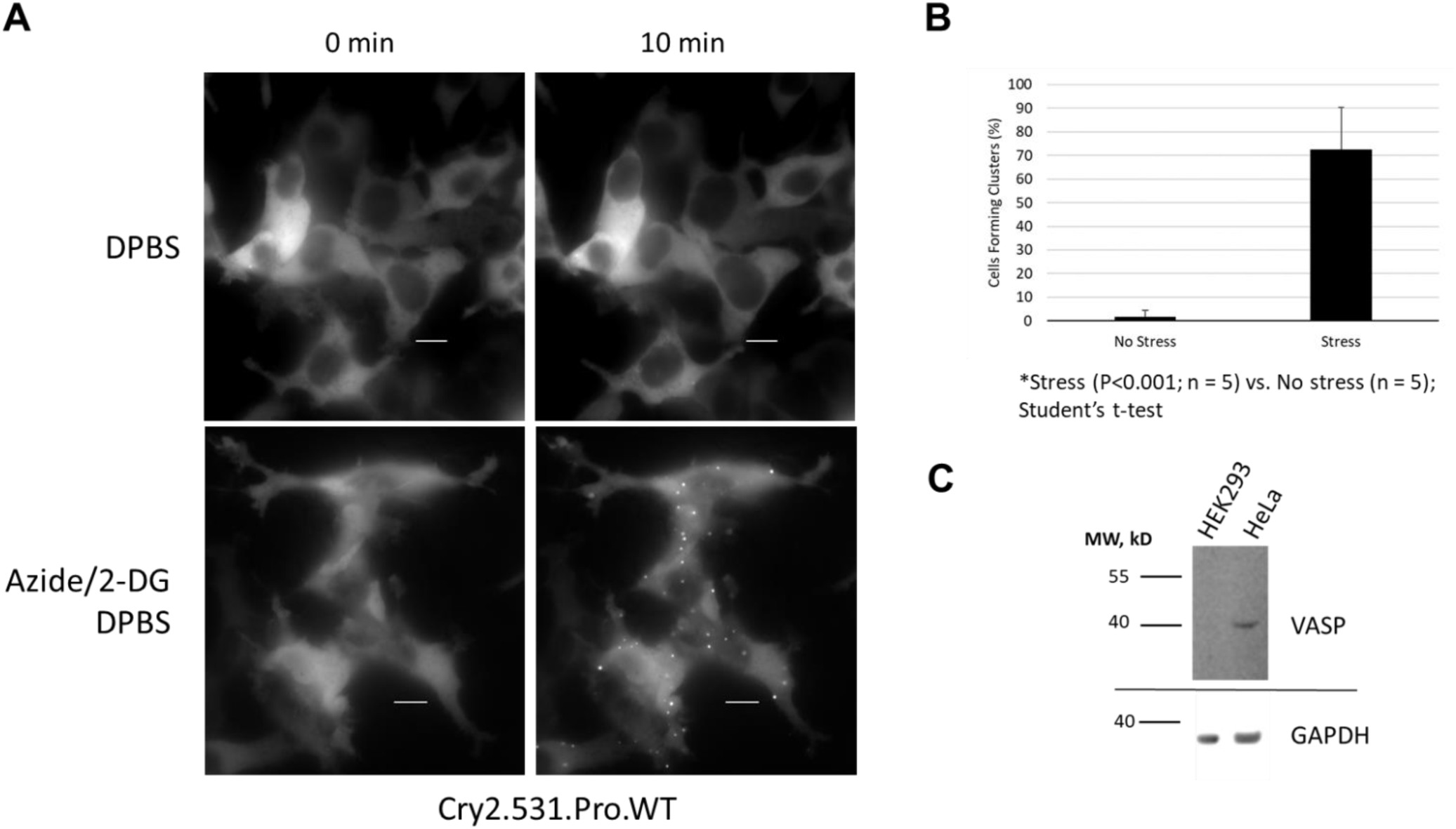
OptoProfilin clustering response in HEK 293T cells. A. OptoProfilin exhibits a clustering response in ATP-depleted 293T cells but not in non-ATP depleted 293T cells. In contrast to HeLa cells, a filopodia recruitment phenotype is absent in non-ATP depleted cells. B. Quantification of experiment described in Panel A. C. Western blot of endogenous VASP levels in HeLa and HEK 293T cells. Scale bars = 10 microns.

**FIGURE 9.**
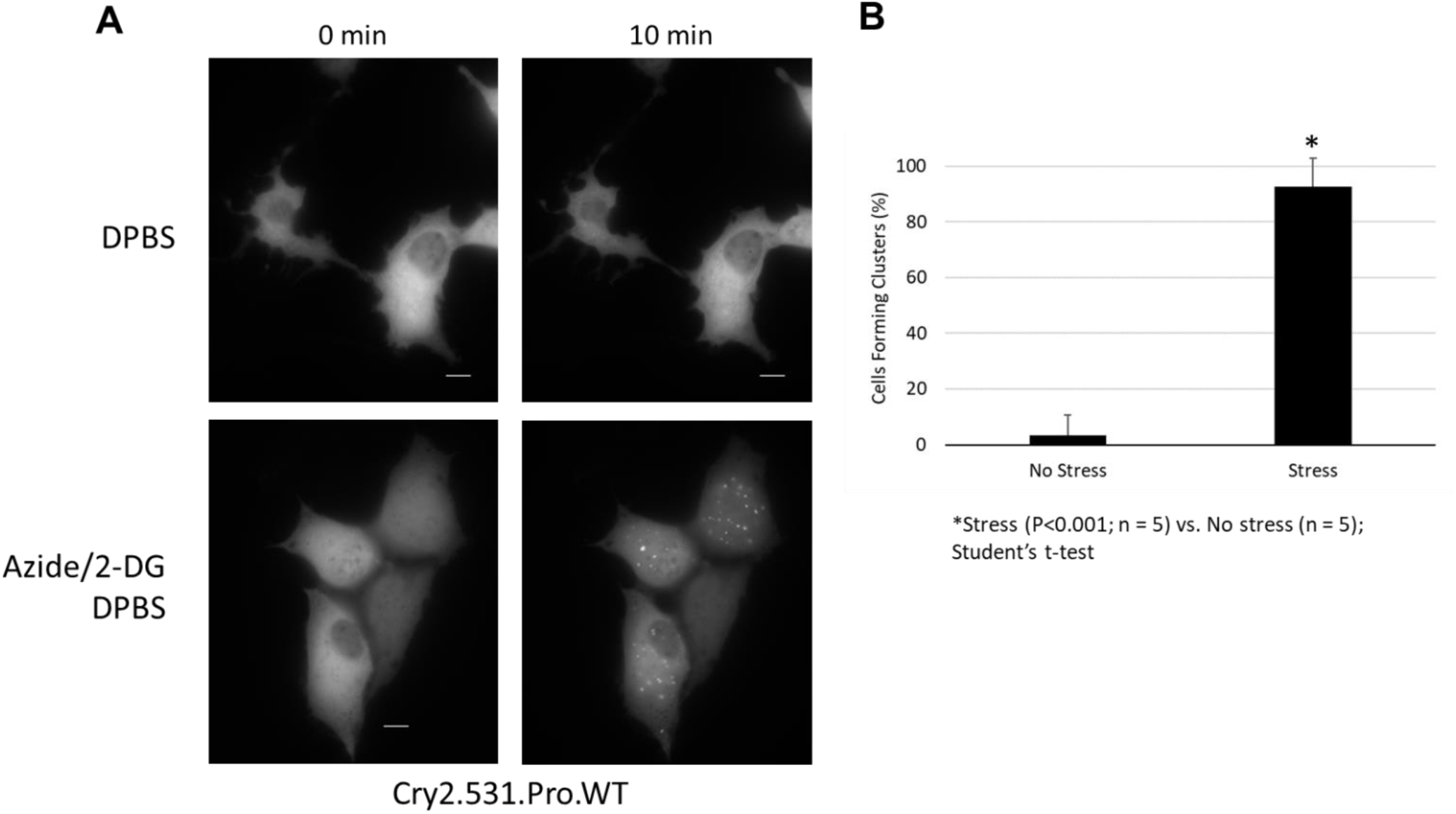
OptoProfilin clustering response in NIH 3T3 cells. A. OptoProfilin exhibits a clustering response in ATP-depleted 3T3 cells but not in non-ATP depleted 3T3 cells. In contrast to HeLa cells, but similar to HEK 293T cells, a filopodia recruitment phenotype is absent in non-ATP depleted cells. B. Quantification of experiment described in Panel A. Scale bars = 10 microns.

**FIGURE 10.**
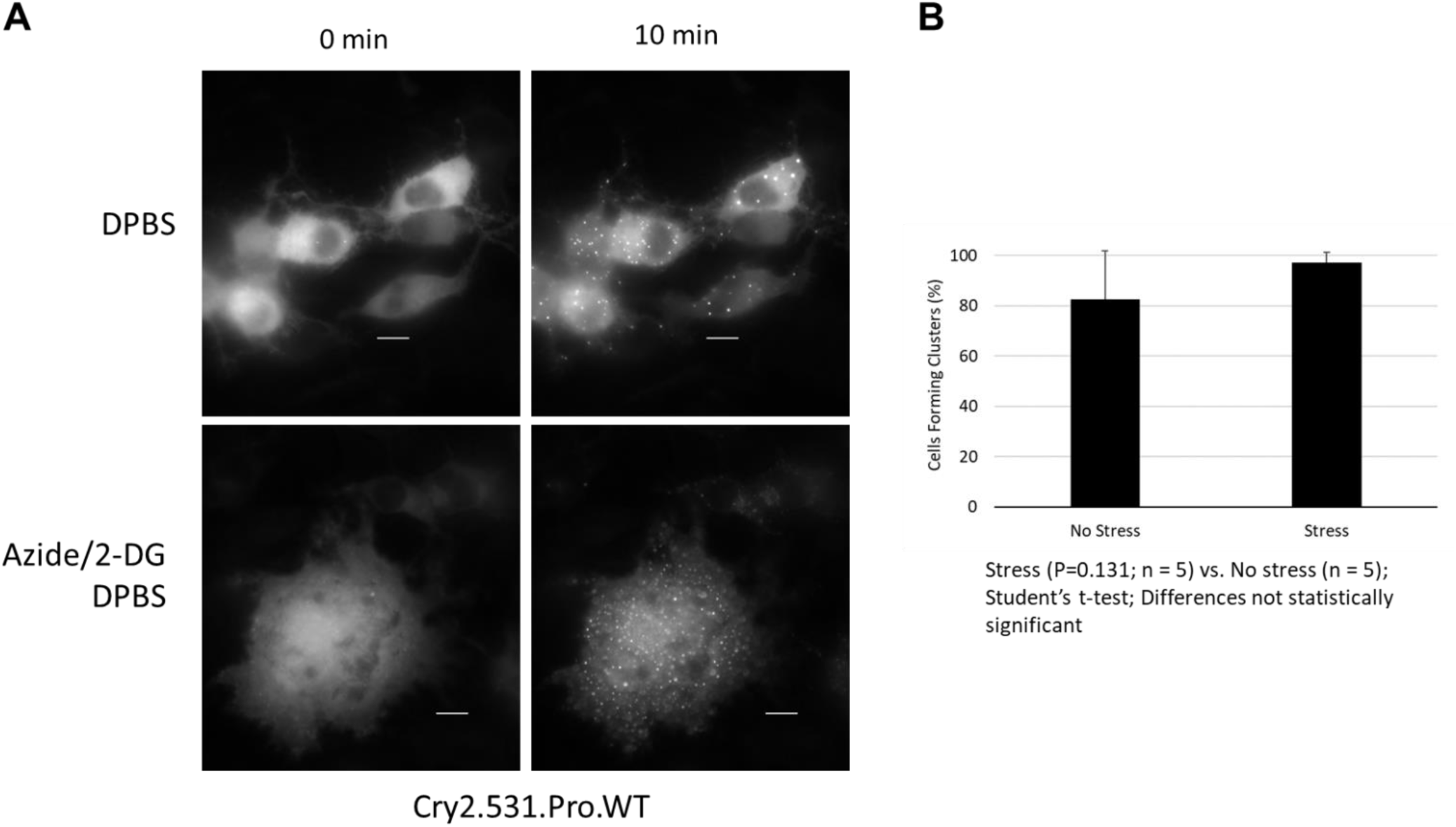
OptoProfilin clustering response in N2a cells. A. OptoProfilin exhibits a clustering response in both ATP-depleted cells and non-ATP depleted cells, in contrast to both HeLa and HEK 293T cells. B. Quantification of experiment described in Panel A. Scale bars = 10 microns.

**FIGURE 11.**
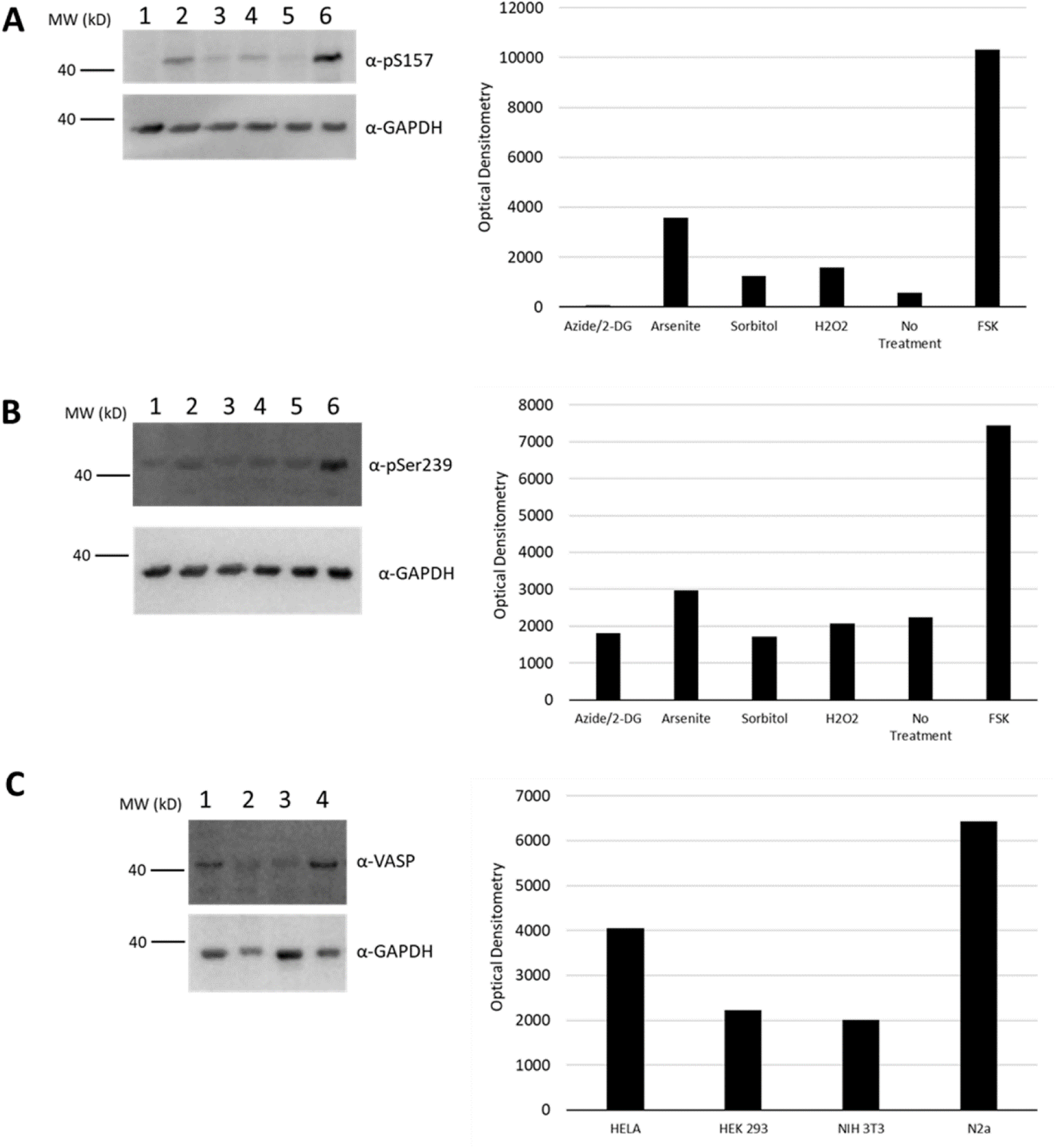
VASP phosphorylation levels in response to various cell treatments and VASP expression across cell lines. A. Western blot of phosphorylation of VASP at Serine 157 in HeLa cells after 15 minutes of the indicated stress treatments (1. 10 mM Azide/6 mM 2-deoxyglucose; 2. 0.5 mM Sodium arsenite; 3. 200 mM Sorbitol; 4. 200 µM H_2_O_2_; 5. No treatment; and 6. 20 µM forskolin). Bar graph shows optical densitometry of bands normalized to GAPDH. B. Phosphorylation of VASP at Serine 239 in HeLa cells after 15 minutes of the indicated stress treatments (same as in panel A). Bar graph shows optical densitometry of bands normalized to GAPDH. C. Western blot of total VASP expression in HeLa, HEK 293, NIH 3T3, and Neuro-2a cell lines. Bar graph shows optical densitometry of bands. Each lane was loaded with 23 µg of cell lysate.

Our proposed mechanism for OptoProfilin function in HeLa cells (**Fig. 12**) anticipates that, under conditions of homeostasis (**Fig. 12A**), light activation and subsequent Cry2-dependent homooligomerization of OptoProfilin results in filopodial recruitment through a multimerization-associated increase in avidity for VASP, which is already present at focal adhesions. While dynamics and stoichiometry of the VASP-Profilin interaction are not completely understood (Hansen and Mullins, 2010), it is reasonable to anticipate that profilin oligomerization could enhance its VASP binding capabilities, as profilin-actin binding is enhanced by profilin in its tetrameric state (Babich et al., 1996). Under stress conditions, VASP is rapidly converted to compact VASP clusters (**Fig. 12B**), which is consistent with observations made in live cell experiments (ex. **Fig. 5**). Upon OptoProfilin activation, this change in VASP distribution is reflected by the appearance of bright, punctate OptoProfilin clusters. Based on the number of different stress stimuli that enable OptoProfilin cluster formation in HeLa cells, it is likely that VASP clustering is the result of a generic stress response mechanism; investigation of the biochemical changes (i.e., dephosphorylation) that may further enhance profilin-VASP interactions and lead to abundant cluster formation are currently underway. We further hypothesize that levels of endogenous VASP expression may be a good predictor of the OptoProfilin response across various cell line. For example, while we have yet to identify a cell line other than HeLa that displays strong focal adhesion recruitment in the absence of stress, we would expect that such cell lines would have similar levels of VASP expression and, of course, similar patterns of VASP subcellular localization.

**FIGURE 12.**
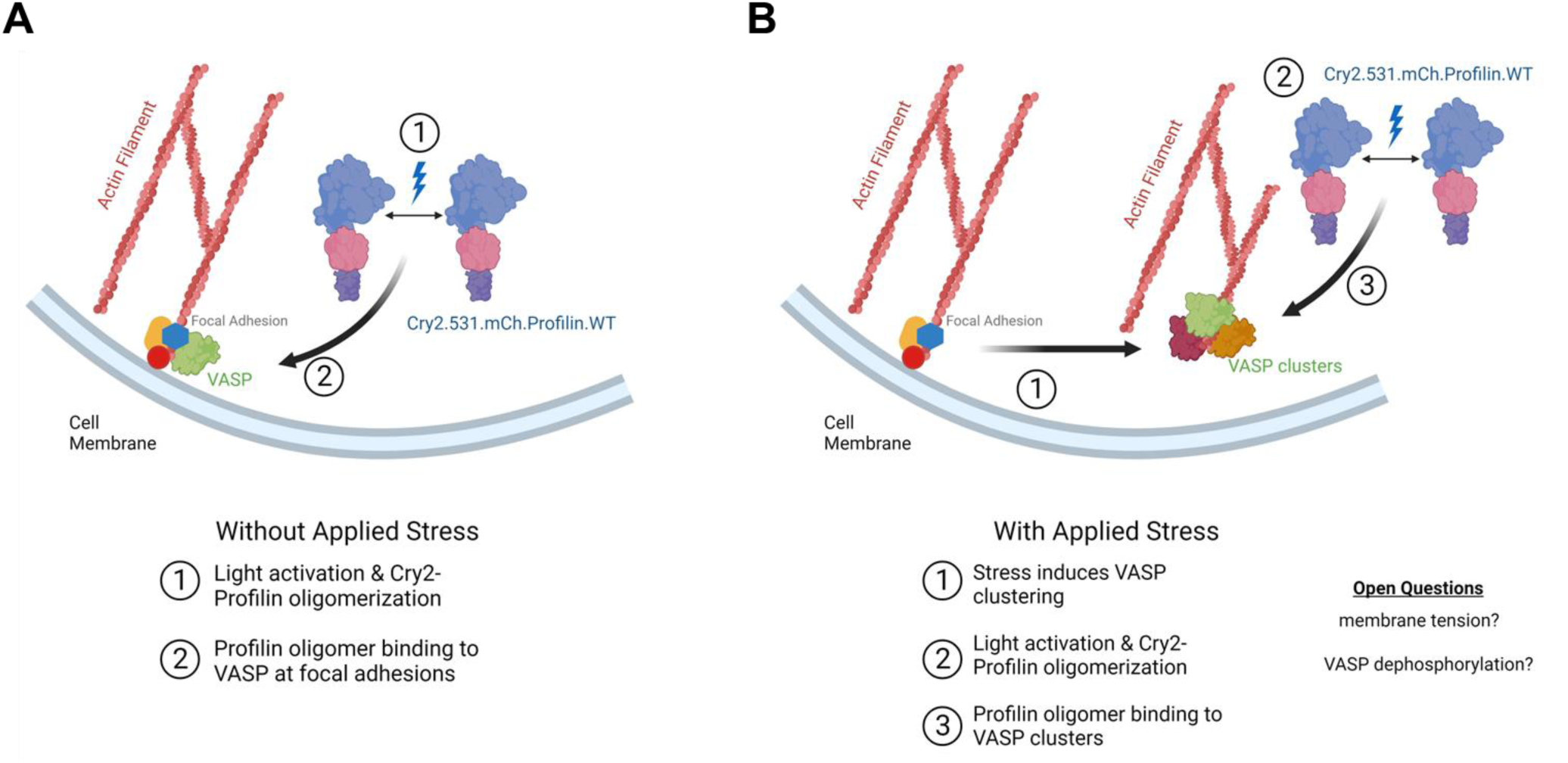
Model of the OptoProfilin light-activated response in HeLa cells. A. Light activated response under non-stressed conditions: Cry2-mediated oligomerization of profilin enhances avidity for VASP-rich filopodia, resulting in abundant filopodial recruitment. B. Light-activated response under stress conditions: applied stress results in enhanced VASP cluster formation. Cry2-mediated oligomerization of profilin initiates recruitment to stress-induced VASP clusters. Changes in phosphorylation state of both VASP and profilin may drive the clustering interaction. The appearance of stress phenotype in response to multiple stress inputs (energetic, oxidative, osmotic) indicates that a general mechanism, such as a change in membrane tension, may underly the observed VASP cluster transition. *Image created with BioRender*.

Future work involving OptoProfilin could proceed in two rather different directions. In the first, the stress response of OptoProfilin could be further explored. While this study employed H_2_O_2_ induced senescence, numerous other reagents (ex. methotrexate; gemcitabine) have been used elsewhere to induce senescence. The capacity of OptoProfilin to detect these small molecule-induced changes in cells could be further investigated. These studies could define whether an OptoProfilin-expressing cell line might have utility as a cellular stress beacon in long term co-culture experiments or in organoids. Additional focus could be placed on the composition of OptoProfilin-VASP clusters themselves. For example, we observed incorporation of GFP-actin (**Supporting Figures 2 and 3**) under ATP-depletion and oxidative stress; other proteins could also be present in these clusters. Elucidation of their identity and relative abundance would further enhance knowledge of the stress-induced cytoskeletal interactome. Such investigations could be bolstered by the application of advanced microscopy techniques, including total internal reflection fluorescence (TIRF) microscopy. In initial TIRF experiments, OptoProfilin was amenable to TIRF imaging under both non-stress and stress conditions (**Supporting Movie 6 and 7**). The second direction could focus on the non-stress associated recruitment to focal adhesions observed in HeLa cells. In these experiments, the ability of OptoProfilin (WT and mutants) to impact cell motility in cancer cells could be investigated. These experiments would define whether OptoProfilin is able to impact cell migration and invasiveness. Finally, while we have used the Cry2 optogenetic system exclusively in this study, it is unknown whether other optogenetic dimerizing/clustering systems could induce the same responses observed with Cry2, or whether these are unique responses that require the Cry2 system. Such benchmarking studies would be instructive for future attempts to design optical biosensors of applied cellular stress.

## MATERIALS AND METHODS

### Plasmids and Cloning

Cloning of Cry2(1-531).mCh.Profilin was conducted using previously reported methods (Wurz et al., 2022a). Briefly, *M. musculus* Profilin (Addgene #56438) was PCR amplified with primers encoding NotI and BsrGI restriction sites. After digestion, these fragments were ligated into a vector containing Cry2(1-531).mCherry to give Cry2(1-531).mCherry.Profilin.WT. Point mutations were introduced into Profilin using mutagenic primers (IDT, Coralville, IA) following a standard site directed mutagenesis protocol using AccuPrime Pfx polymerase (Invitrogen). The actin encoding gene (pCAG-mGFP-actin; Addgene #21948) construct was a generous gift from Ryohei Yasuda (Murakoshi et al., 2008). EGFP-Profilin-10 was a gift from Michael Davidson (Addgene plasmid # 56438 ; http://n2t.net/addgene:56438 ; RRID:Addgene_56438)

### Cell lines and transfection

Midi prep quantities of DNA of each construct were generated from *E. coli* stocks and purified for cell transfection. Transfection of cells (HeLa, HEK 293T, NIH 3T3, and N2a) was then performed with the Calfectin reagent (SignaGen) following manufacturer’s suggested protocols. Briefly, for dual transfections in 35 mm glass bottom dishes, plasmid DNA was combined in a 1:1 ratio (1,000 ng per plasmid) in 100 ul of DMEM (without serum), followed by the addition of 3 ul of Calfectin reagent. Transfection complexes were incubated no longer than 15 minutes at room temperature prior to adding to cells. For single transfections in 35 mm glass bottom dishes, 1,000 ng of plasmid DNA was used per transfection. Transfection solutions were allowed to remain on cells overnight. Cells were maintained at 37°C and 5% CO_2_ in a humidified tissue culture incubator, in culture medium consisting of DMEM supplemented with 10% FBS and 1% Penicillin-Streptomycin.

### Cell treatments, Imaging, and Immunofluorescence

#### Live cell experiments

Transfected cells (Hela, HEK 293T, NIH 3T3, N2a) were washed Dulbecco’s PBS (with calcium and magnesium; 1 x 1 mL), prior to treatment with cell stress media (ATP depletion medium (6 mM D-Deoxyglucose and 10 mM Sodium Azide in DPBS); Oxidative stress media (0.5 mM Sodium Arsenite in DPBS); Osmotic stress media (200 mM Sorbitol in DPBS); or Senescence media (200 uM H2O2 in DPBS, 2 h, followed by washing with DPBS and restoration of DMEM with 10% FBS). Cells were allowed to equilibrate in the live cell incubation system (OKOLab) for 15 min prior to beginning the illumination sequence. The unstressed to stress transition experiment in HeLa cells was performed by adding a 200 uL aliquot of 100 mM NaN3/60 mM 2-DG to the cells in 2 mL of DPBS during the experiment to give a final concentration of 6 mM D-Deoxyglucose and 10 mM Sodium Azide.

#### Fixed cell experiments

Transfected HeLa cells were washed with Dulbecco’s PBS (with calcium and magnesium; 3 x 1 mL), prior to treatment with ATP depletion medium (6 mM D-Deoxyglucose and 10 mM Sodium Azide in Dulbecco’s PBS) or Sodium Arsenite (0.5 mM in DPBS). Cells were returned to the 37 °C incubator for 10 min, then removed and exposed to blue light (Sunlite LED Par30 Reflector, Item #80021, 4 Watts, 120 Volt, 66.21 µmol/s/m^2^; placed 10 cm from cell culture dishes) for 2 min. The stress medium was removed by aspiration, cells washed gently (1X with 1 mL Dulbecco’s PBS), then fixed for 10 min with pre-warmed 4% Paraformaldehyde solution (37°C; prepared from 16% PFA (Electron Microscopy Sciences and DPBS) at 37°C. Following removal of fixative solution, cells were washed with PBS, then permeabilized for 30 min using antibody dilution buffer (30 µl Triton X-100, 0.1 g of BSA, 10 mL of Dulbecco’s PBS). Cells then were incubated overnight at 4°C with primary antibody (anti-VASP (Cell Signaling Technologies), anti-Paxillin (Cell Signaling Technologies), or anti-Ataxin2 (ProteinTech); 1:500 in antibody dilution buffer). The following day, primary antibody solution was removed by pipette, and cells were washed three times with Dulbecco’s PBS. Cells were incubated with Alexa 488 conjugated goat anti-rabbit secondary (Invitrogen; 1:1000 in antibody dilution buffer) for 1 hour at room temperature, followed by a Dulbecco’s PBS wash (1 mL; 3 x 5 min). Cells were stored in Dulbecco’s PBS prior to imaging.

##### Confocal Microscopy

Confocal images of fixed cells were collected with a Zeiss LSM 700 laser scanning microscope using ZEN Black 2012 software. Fluorescence images were colorized and overlaid using FIJI software.

##### Widefield Microscopy

A Leica DMi8 Live Cell Imaging System, equipped with an OKOLab stage-top live cell incubation system, LASX software, Leica HCX PL APO 63x/1.40-0.60na oil objective, Lumencor LED light engine, CTRadvanced+ power supply, and a Leica DFC900 GT camera, was used to acquire images. Exposure times were set at 50 ms (GFP, 470 nm) and 200 ms (mCherry, 550 nm), with LED light sources at 50% power, and images acquired every 30 seconds over a 10 min time course. TIRF imaging was conducted using a 100X TIRF compatible objective and 10% laser power.

### Western blotting

HEK 293T cells were lysed 16 hours post-transfection with 200 µL of M-PER lysis buffer (Thermo Scientific) plus protease inhibitors. After 10 min on a rotary shaker at room temperature, lysates were collected and centrifuged for 15 min (94 rcf; 4°C). The supernatants were collected and combined with 6X Laemmli sample buffer, followed by heating for 10 min at 65°C. The resulting samples were subjected to electrophoresis on a 10% SDS-PAGE gel and then transferred onto PVDF membranes (20 V, overnight, at 4 °C). Membranes were then blocked for 1 h with 5% BSA in TBS with 1% Tween (TBST), followed by incubation with primary antibody (Anti-mCherry antibody (Cell Signaling); 1:1000 dilution in 5% BSA – TBST; Anti-GAPDH antibody (ThermoScientific); 1:1000 dilution in 5% BSA – TBST) overnight at 4°C on a platform rocker. The membranes were then washed 3 x 5 min each with TBST and incubated with the appropriate secondary antibody in 5% BSA – TBST for 2 hours at room temp. After washing 3 x 5 min with TBST, the membranes were exposed to a chemiluminescent substrate for 5 min and imaged using an Azure imaging station.

## ACKNOWLEDGEMENTS

We acknowledge financial support from the NIH (1R15NS125564-01) to RMH and the ECU Brody Summer Biomedical Research Program (KW).

## SUPPORTING INFORMATION

**Supporting Figure 1.**
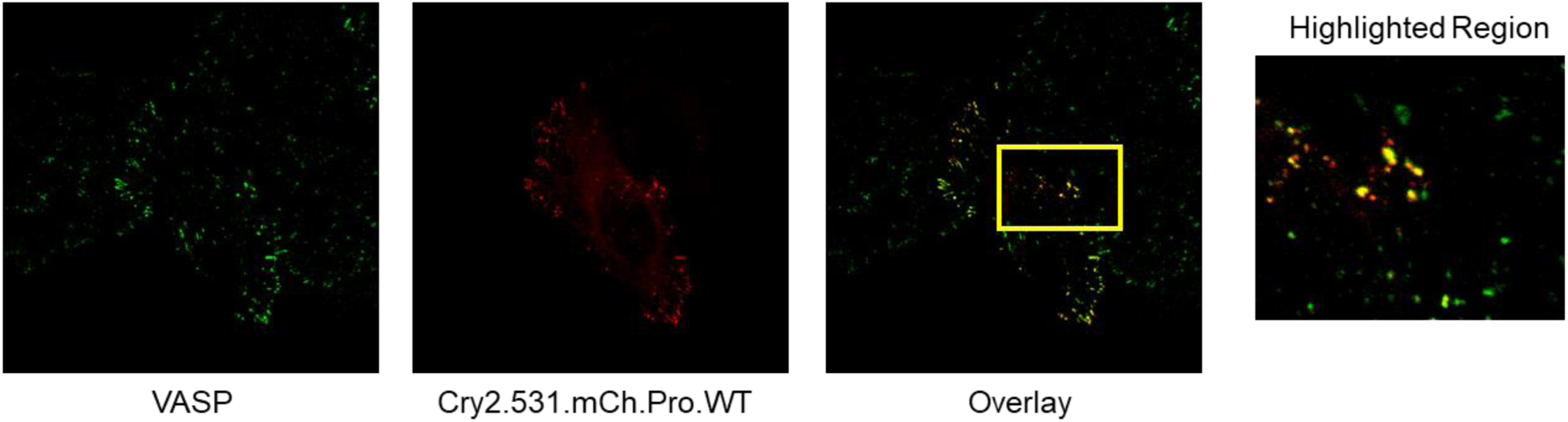
Immunofluorescence of VASP-rich clusters formed in sodium arsenite-treated HeLa cells. HeLa cells were treated with 0.5 mM Sodium Arsenite in DPBS for 15 minutes prior to fixation (4% PFA, 10 min at 37 °C). Immunostaining of VASP (green, Alexa 488 secondary antibody) reveals that OptoProfilin clusters (mCherry, red) formed in response to light activation are VASP-rich.

**Supporting Figure 2.**
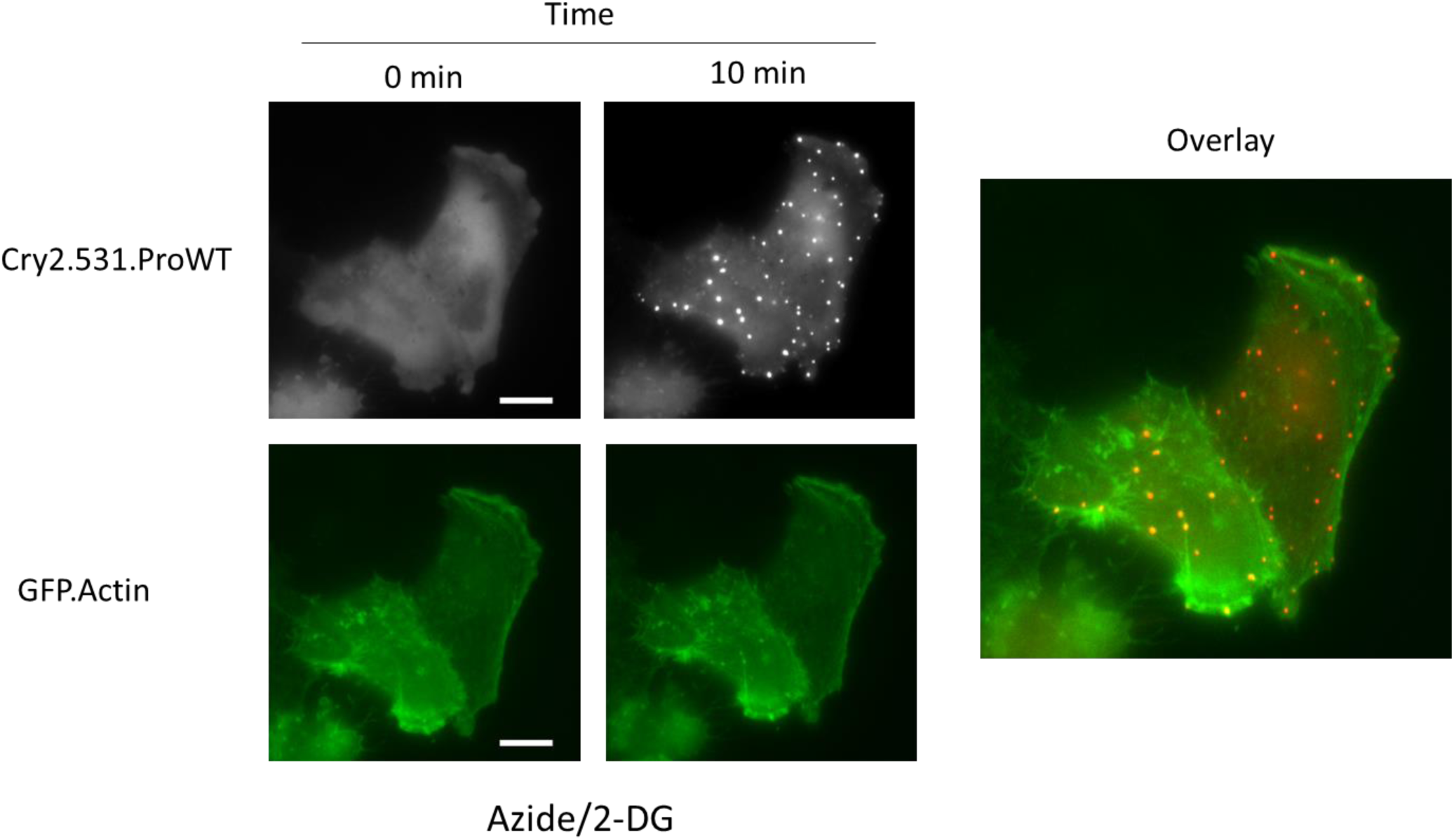
Actin is incorporated into OptoProfilin clusters formed under ATP-depletion conditions. Live-cell imaging of ATP-depleted HeLa cells co-transfected with OptoProfilin and a GFP-Actin expression construct reveal that actin is incorporated into OptoProfilin clusters. Cells were treated with ATP-depletion media (10 mM NaN3/6 mM 2-DG/DPBS) for 15 minutes prior to a 10 minute timecourse of light activation (470 nm, 50 ms pulse, applied every 30 s) and imaging (mCherry and GFP channels).

**Supporting Figure 3.**
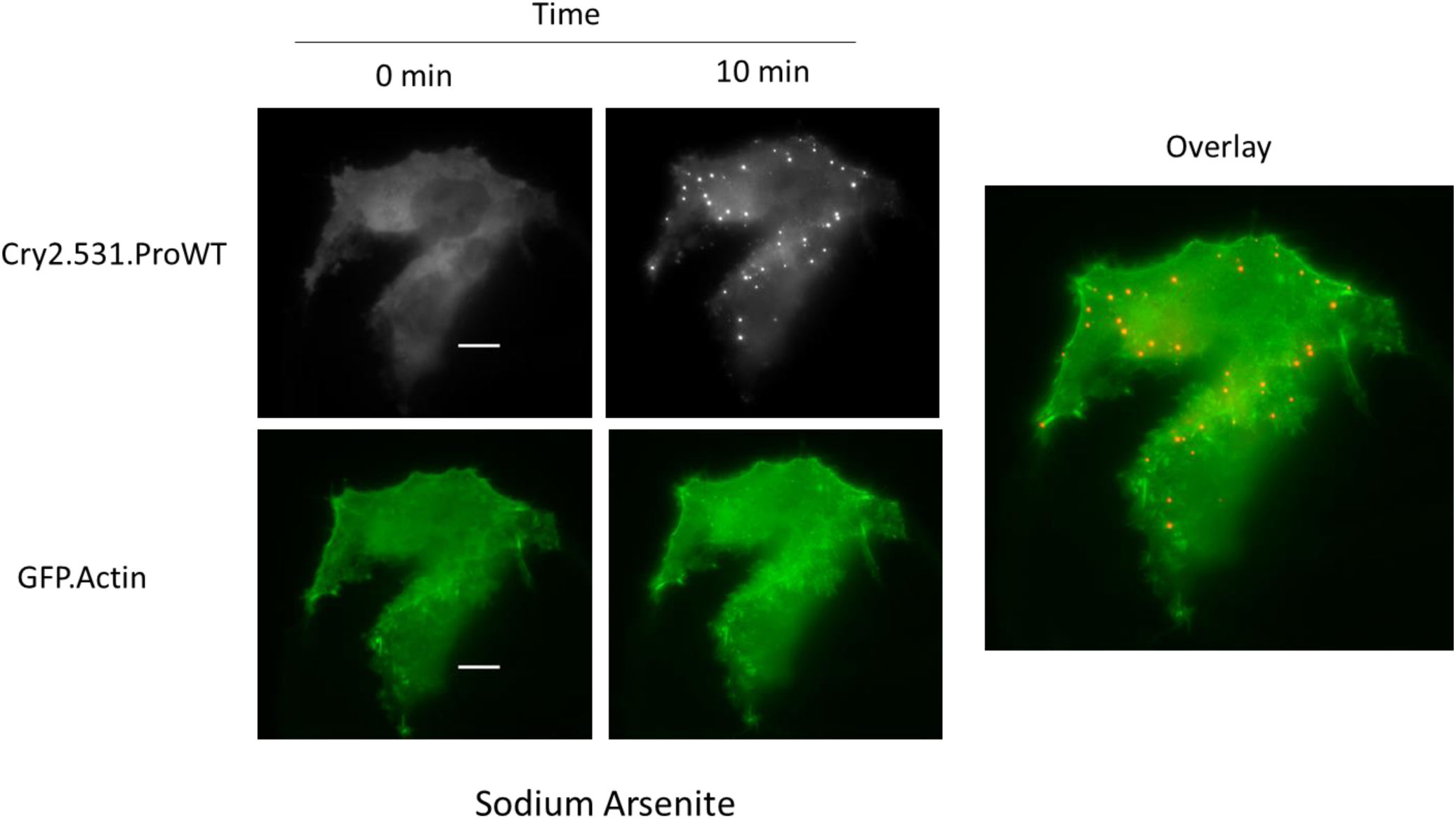
Actin is incorporated into OptoProfilin clusters formed under oxidative stress conditions. Live-cell imaging of oxidative stress treated HeLa cells co-transfected with OptoProfilin and a GFP-Actin expression construct reveal that actin is incorporated into OptoProfilin clusters. Cells were treated with ATP-depletion media (0.5 mM Sodium Arsenite/DPBS) for 15 minutes prior to a 10 minute timecourse of light activation (470 nm, 50 ms pulse, applied every 30 s) and imaging (mCherry and GFP channels).

