## Supplementary figures and images for "OptoProfilin: A Single Component Biosensor of Applied Cellular Stress"

### SUPPORTING MOVIE 1

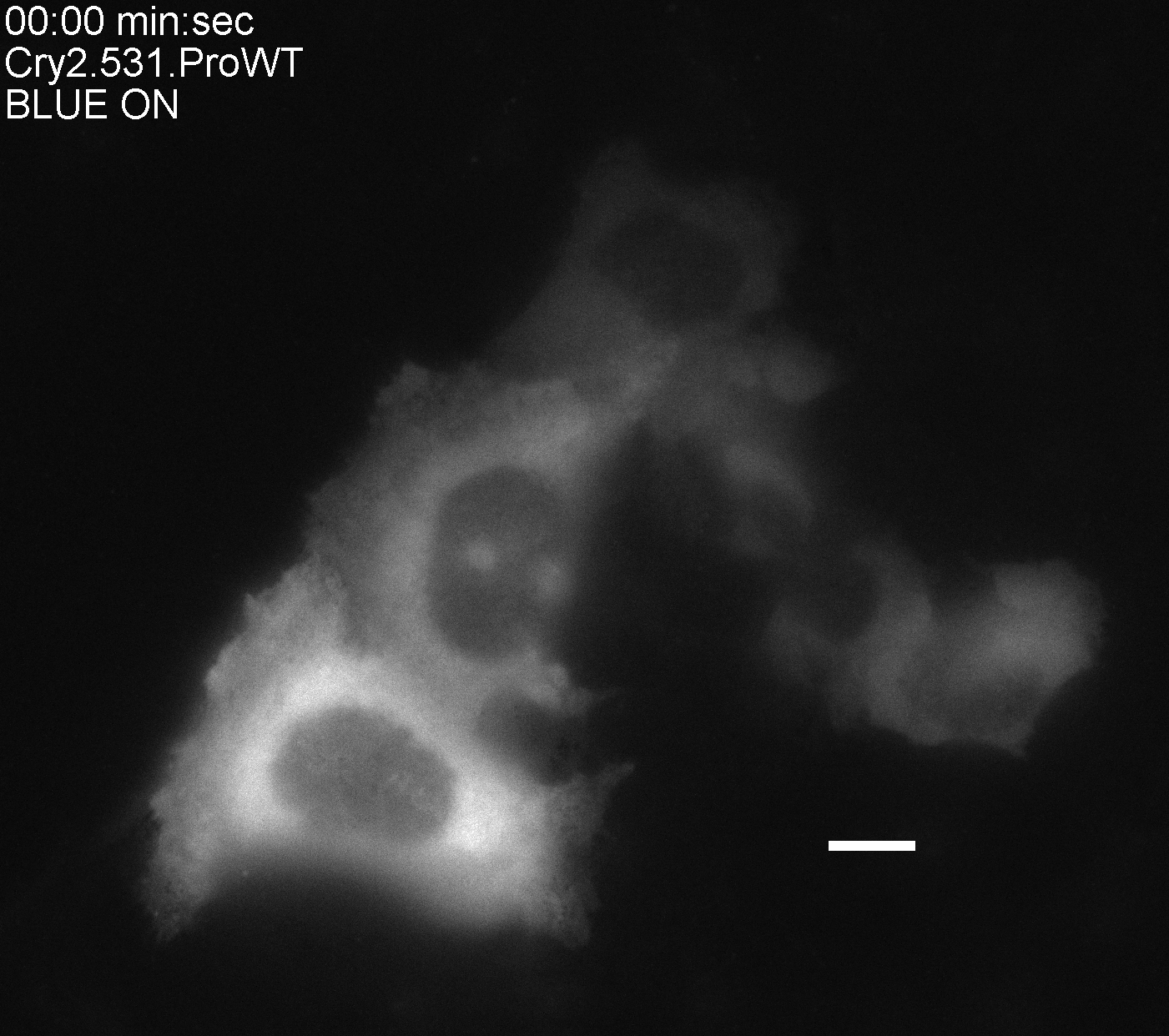

### SUPPORTING MOVIE 2

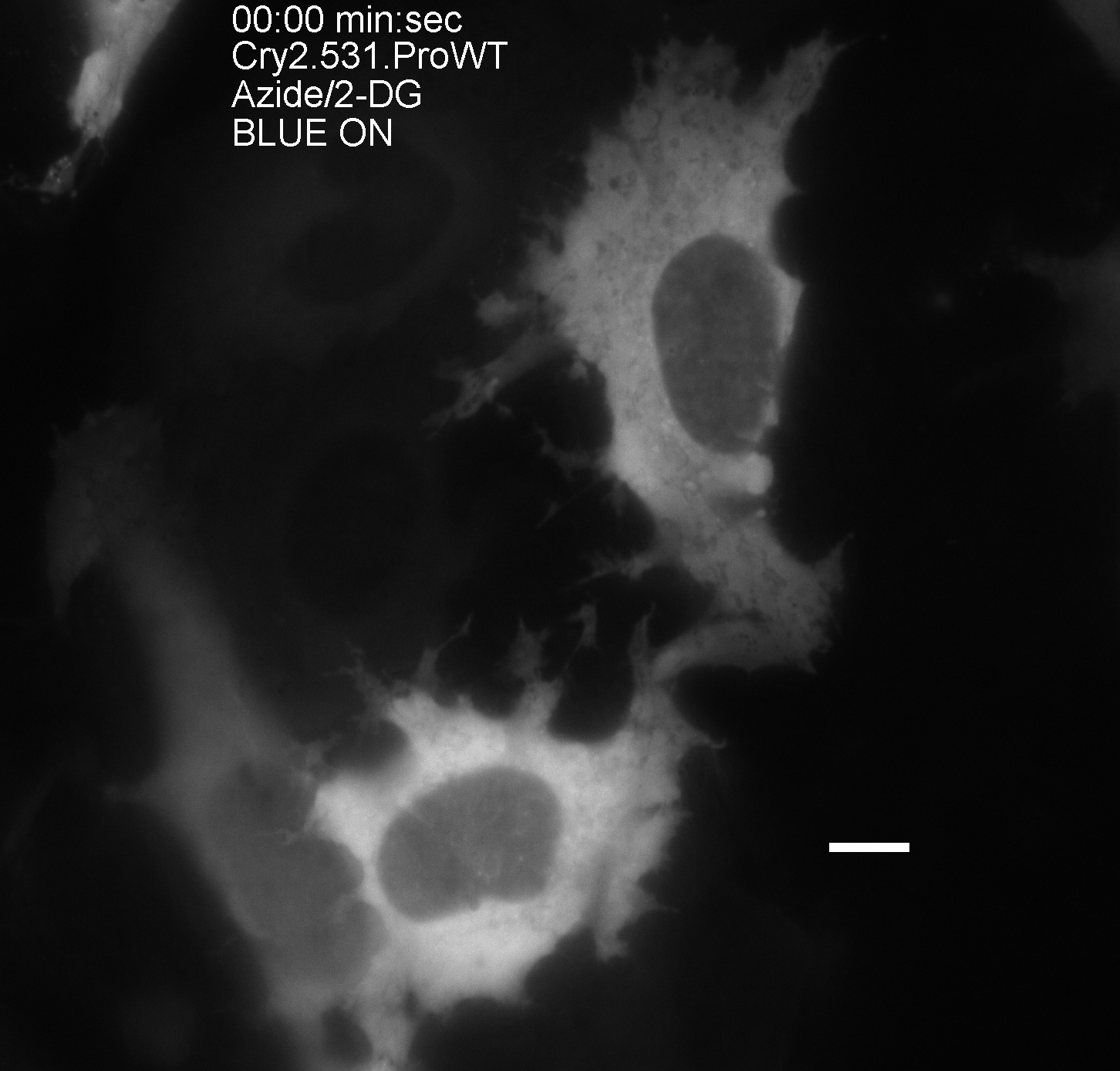

### SUPPORTING MOVIE 3

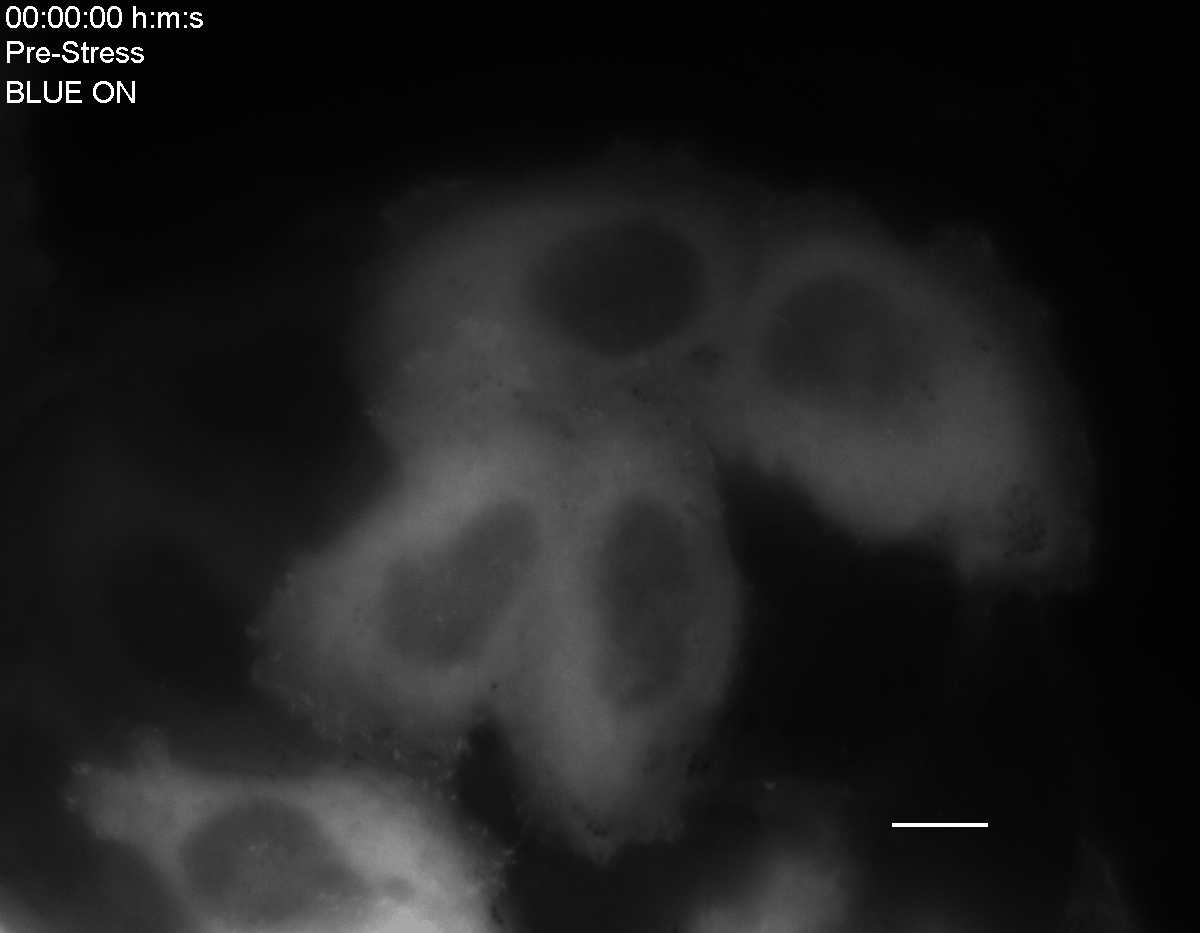

### SUPPORTING MOVIE 4

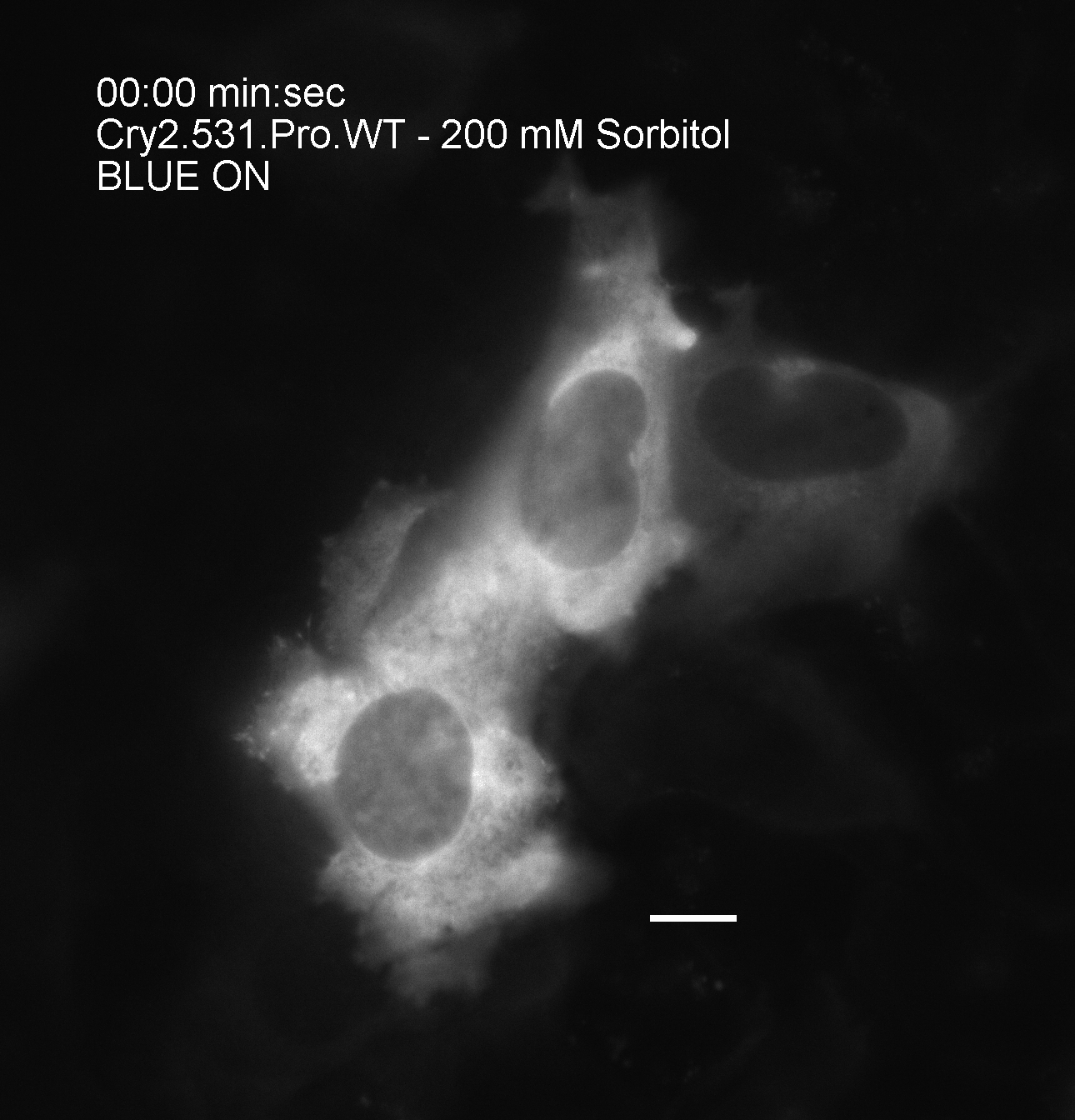

### SUPPORTING MOVIE 5

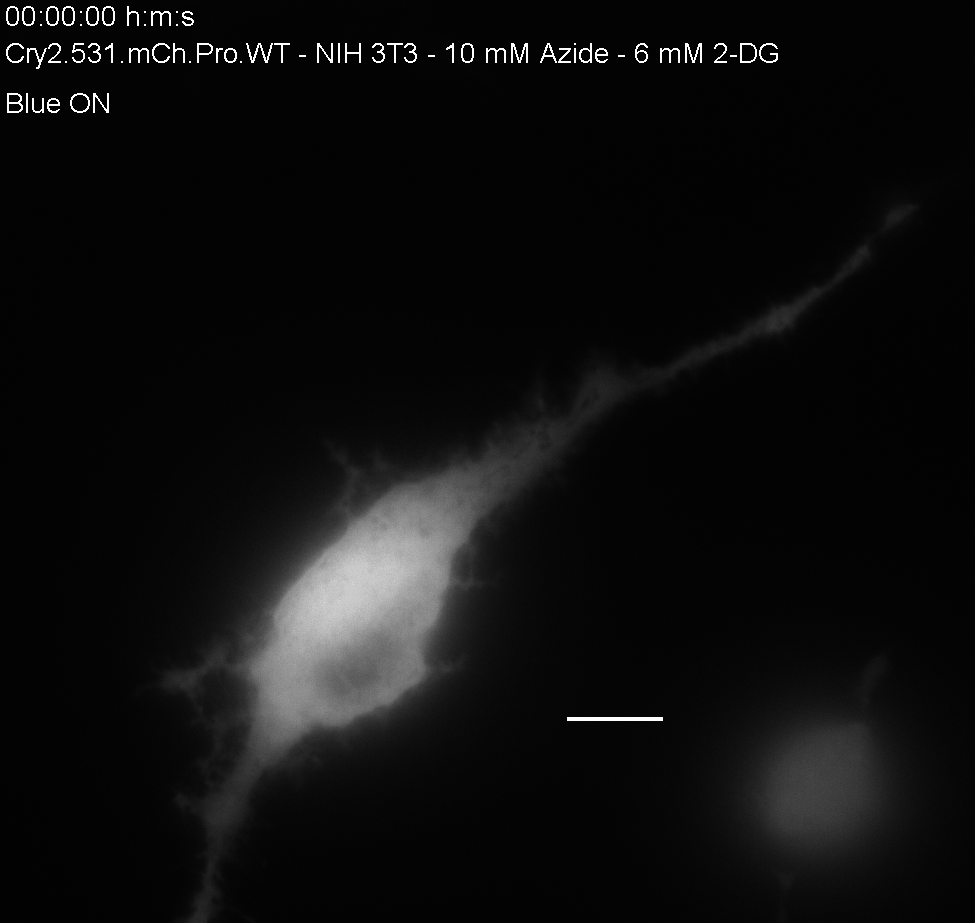

### SUPPORTING MOVIE 6

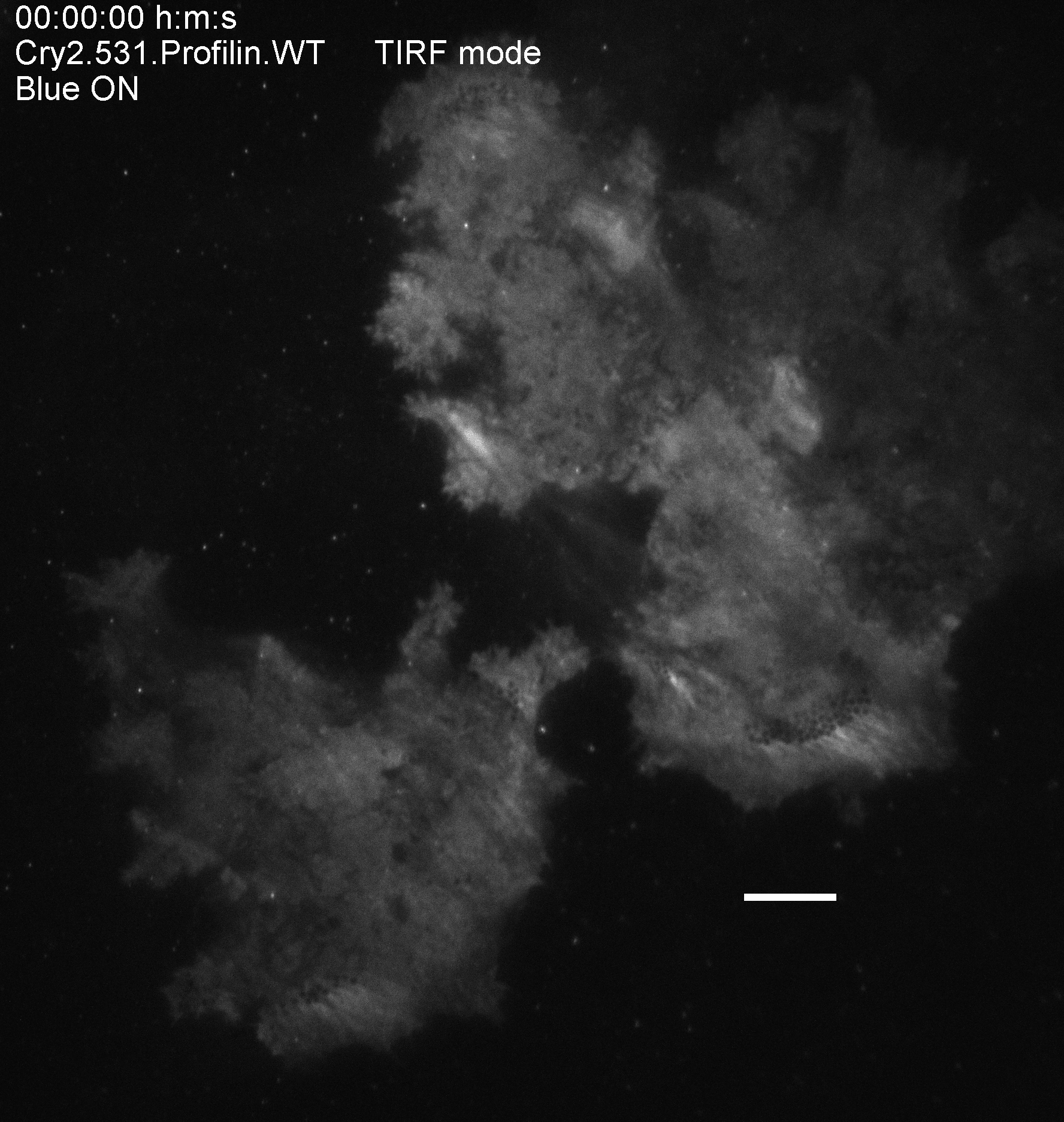

### SUPPORTING MOVIE 7

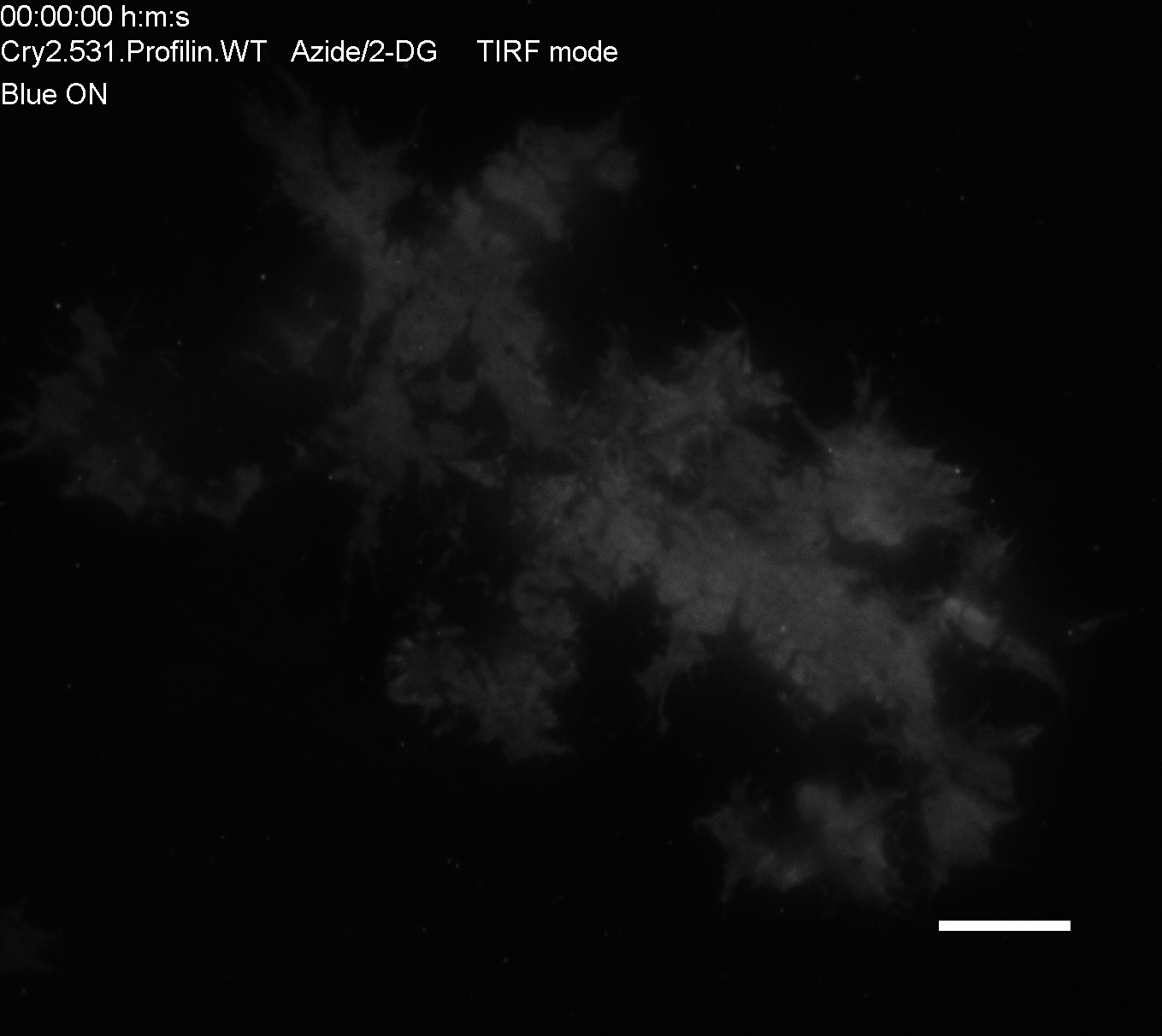
